# A validated generally applicable approach using the systematic assessment of disease modules by GWAS reveals a multi-omic module strongly associated with risk factors in multiple sclerosis

**DOI:** 10.1101/2020.10.26.351783

**Authors:** Tejaswi V.S. Badam, Hendrik A. de Weerd, David Martínez-Enguita, Tomas Olsson, Lars Alfredsson, Ingrid Kockum, Maja Jagodic, Zelmina Lubovac-Pilav, Mika Gustafsson

**Affiliations:** School of Bioscience, Systems Biology Research Center, University of Skövde, Sweden; Bioinformatics, Department of Physics, Chemistry and Biology, Linköping university, Linköping, Sweden; Department of Clinical Neuroscience, Karolinska Institutet, Center for Molecular Medicine, Karolinska University Hospital, SE-171 76, Stockholm, Sweden; Institute of Environmental Medicine, Karolinska Institutet, Center for Molecular Medicine, Karolinska University Hospital, SE-171 76, Stockholm, Sweden

**Author notes:** Corresponding author: Mika Gustafsson. These authors contributed equally to the work (Tejaswi V.S. Badam, Hendrik A. de Weerd). These authors share senior authorship (Zelmina Lubovac-Pilav, Mika Gustafsson). **SUMMARY**: Our benchmark of multi-omic modules and validated translational systems medicine workflow for dissecting complex diseases resulted in multi-omic module of 220 genes highly enriched for risk factors associated with multiple sclerosis.

**Keywords:** Benchmark, Multi-omics, Network modules, Multiple Sclerosis, Risk factors

## Abstract

**Background:** There are few (if any) practical guidelines for predictive and falsifiable multi-omics data integration that systematically integrate existing knowledge. Disease modules are popular concepts for interpreting genome-wide studies in medicine but have so far not been systematically evaluated and may lead to corroborating multi-omic modules.

**Methods:** We assessed eight module identification methods in 57 previously published expression and methylation studies of 19 diseases using GWAS enrichment analysis. Next, we applied the same strategy for multi-omics integration of 19 datasets of multiple sclerosis (MS), and further validated the resulting module using both GWAS and risk-factor associated genes from several independent cohorts.

**Results:** Our benchmark of modules showed that in immune-associated diseases modules inferred from clique-based methods were the most enriched for GWAS-genes. The multi-omics case study using MS revealed the robust identification of a module of 220 genes. Strikingly, most genes of the module was differentially methylated upon the action of one or several environmental risk factors in MS (n = 217, P = 10^-47^) and were also independently validated for association with five different risk factors of MS, which further stressed the high genetic and epigenetic relevance of the module for MS.

**Conclusion:** We believe our analysis provides a workflow for selecting modules and our benchmark study may help further improvement of disease module methods. Moreover, we also stress that our methodology is generally applicable for combining and assessing the performance of multi-omics approaches for complex diseases.

## INTRODUCTION

Complex diseases are the result of disruptions of many interconnected multimolecular pathways, reflected in multiple omics layers of regulation of cellular function, rather than perturbations of a single gene or protein[1]. Systems and network medicine aim to translate observed omics differences in patients using networks, in order to personalize medicine[2]. Importantly, genes that are associated with diseases are more likely to interact with each other rather than with non-disease associated genes, forming multi-omics network disease modules[3,4]. Owing to the incompleteness of the underlying multi-omics interactions, the networks are often modeled as effective gene-gene interactions, using for example STRING database[5]. Thus, network modules might be ideal tools for multi-omics analysis. However, the evaluation of performance of different module inference methods remains a poorly understood topic, which creates the need for transparent evaluation of these methods based on objective benchmarks across various diseases and omics. Genomic concordance has been suggested as a multi-omics validation principle[4,6], i.e., modules derived from one omic, such as gene expression or DNA methylation should be enriched for disease-associated single nucleotide polymorphisms (SNPs).

The variety of algorithms that have been proposed and applied for identification of disease modules can be categorized into two main groups. On the one hand, there are methods which rely purely on clustering of the genes in relevant disease networks[7]. On the other hand, there are algorithms which make use of disease-associated molecules or genetic loci to reveal disease modules that correlate with disease function, such as the disease module detection (DIAMOnD) algorithm[8], clique-based methods[9],[10] and weighted gene co-expression network analysis (WGCNA)[11]. The data-derived information can either be differentially expressed genes or differentially correlated or co-expressed genes. Methods following the former approach were recently benchmarked by a metric utilizing genomic concordance within the DREAM consortia[12]. However, so far, algorithms from the latter group have not been benchmarked.

In this study we analyzed, assessed, and compared the performance of eight of the most popular methods for disease module analysis using the R package MODifieR[13] on 19 different diseases including 47 expression and ten methylation datasets. We assessed the performance of the methods using genome-wide association (GWAS) enrichment analysis from the summary statistics of all assayed SNPs similarly as in DREAM[12]. The resulting workflow provided a systematic procedure for selecting the best method for each disease and set the stage for method development in the disease module area. Moreover, it allowed the predictive assessment of combining multiple datasets across several omics using GWAS, which we tested in multiple sclerosis (MS), a heterogeneous complex disease. Briefly, we derived multi-omic modules in a stepwise optimization of GWAS enrichment from transcriptomic and methylomic analyses of MS. We further evaluated the identified multi-omic MS module of 220 genes for its enrichment across DNA methylation studies of eight known lifestyle-associated risk factors of MS. Additionally, we validated the identified significant enrichment risk factors in an independent DNA methylation MS study which indeed showed a very strong and significant MS enrichment for both module genes and risk factor associations. In summary, we provide a robust multi-omics strategy that can be used to disentangle networks of affected genes in complex diseases from both genetic and environmental levels.

## MATERIALS AND METHODS

### Benchmark data

A total of 47 publicly available datasets for the transcriptomic benchmark and ten publicly available datasets for the methylomic benchmark were used. To avoid bias due to subtypes of diseases and drug treatments, we searched for datasets that have only patient and control samples, and that are available for download from the GEO database. We categorized the datasets into seven distinct disease types based on the disease-trait type associations used in Choobdar et al[12]., i.e. autoimmune, cardiovascular, glycemic, inflammatory, neurodegenerative, and psychiatric and social disorders. A total of 19 complex diseases were used in the transcriptomic benchmark analysis, while six complex diseases were used in the methylation benchmark analysis. The methylation benchmark diseases belong to inflammatory, autoimmune, and glycemic disease types.

### MS use case data

A total of 14 publicly available and one non-publicly available transcriptomic and methylomic MS-related datasets were used in the MS multi-omics integration use case. In general, every dataset in the MODifieR benchmark was also used in the MS use case, with exceptions according to certain criteria. The inclusion of transcriptomic MS datasets followed the criteria: 1) The largest dataset by sample number, per tissue, is shown in the MODifieR benchmark; 2) Replication cohorts are not included in the MS use case. Criteria for inclusion of methylomic MS datasets were the following: 1) The largest dataset by sample number, per tissue or cell type, is included in the MODifieR benchmark; 2) A single dataset for every cell-specific tissue was included in the benchmark; 3) Methylation studies that reported using whole blood as sample tissue were excluded from the MS use case, due to the high heterogeneity of this type of data.

For the additional independent validation, we utilized the methylation microarray analysis of 279 blood samples analyzing from Kular *et al* ^25^. For each of these MS patients (n_MS_= 139) and healthy controls (n_HC_= 140), we also collected their lifestyle-associated risk factors from questionnaires that were part of the Epidemiological Investigation of Multiple Sclerosis (EIMS) study. Those factors were smoking status, prior EBV infection, sunbathing, nightshift work, alcohol consumption, as well as phenotypic features (age, sex, BMI at age of 20).

### Pre-processing and quality control of risk factor methylation data

DNA methylation datasets were downloaded from GEO as raw IDAT files, when available, or matrices of beta values. Pre-processing of the data was performed using the Chip Analysis Methylation Pipeline (ChAMP) R package[14], version 2.16.2. Default parameters were used for probe and sample filtering. Probes with a detection P-value above 0.01, probes with a fraction of failed (bead count less than 3) samples over 0.05, non-CpG probes, SNP-related probes, multi-hit probes, and probes located on chromosomes X and Y, were removed. Samples with a proportion of failed (NA) probe P-values over 0.1 were also removed from the analysis. Post-filtering imputation of NA values was conducted on the beta matrices, with default parameters (“combine” method, k = 5, probe cutoff = 0.2, sample cutoff = 0.1). Filtered imputed matrices were normalized applying the Beta-Mixture Quantile dilation (BMIQ) normalization method[15]⍰, including correction of Type-I and Type-II probe effects. Data quality was assessed by producing multi-dimensional scaling (MDS) plots of the top 1,000 most variable positions per sample, density plots for the distribution of beta values, and hierarchical clustering of samples, before and after normalization. Singular value decomposition (SVD) was used to detect the most significant components of variation in the data. Unwanted sources of variation in the normalized data were corrected using ComBat batch effect correction[16].

### Module Identification

The MODifieR^13^ R package offers nine different methods for producing disease modules for which we included all but Clique SuM exact as it is highly similar to Clique SuM. The included methods will produce modules based on the provided omics input and background network and do not include prioritization of pathway association. MODifieR methods used for module identification through this study are listed in the Supplementary Table 3. For the methods that require a network, we used the human PPI network from STRING^5^ database version 11, consisting of 11,295,036 interactions among 18,746 unique genes/proteins. We filtered the network to have high confidence interactions by using the cutoff > 900 to reduce the number of false positives, resulting in a subset of 631,782 interactions between 12,123 unique genes/proteins. For co-expression methods, the network is computed within the method algorithm from the gene expression matrix. In case of the benchmark analysis, we used a stringent cutoff of score > 900, so that the runs were not computationally intensive. For the MS use case benchmark, we used the network combined score cutoff > 700. The processed matrix for each dataset and their respective phenotypic information were downloaded from GEO. The input object is prepared using the *create_input_microarray* function from the MODifieR package which is then used for creating the modules. The input function applies linear model using limma for comparison of patient’s vs controls to get the differentially methylated or expressed genes. A dynamic cutoff of 5% in the differentially methylated or expressed genes is applied for input seed genes for the methods that require seed genes.

### Differential methylation analysis of risk factor data

Differentially methylated probes (DMPs) were found by fitting a linear model to the data using the limma R package[17]⍰, version 3.42.2 implemented in the ChAMP function *champ.DMP*. P-values were adjusted for multiple testing using Benjamini-Hochberg False Discovery Rate (FDR) correction. Differentially methylated genes (DMGs) were obtained and annotated using the org.Hs.eg.db R package⍰, version 3.10.0. DMG lists were cross-checked against the STRING database version 11 PPI network used for module identification in the MS multi-omics approach (high confidence interactions, combined score > 700). DMGs that were not present in the PPI network were removed. In case of the additional MS validation dataset, a linear mixed effect model with risk factors (age, sex, BMI at age of 20, smoking, alcohol consumption, sun exposure, night shift work, contact with organic solvents) as categorical covariates was implemented to find the differentially methylated genes after the preprocessing step, as described in the preprocessing section of the methods. Since all the patients were EBV positive, we did not include it for linear mixed effect model.

### Validation of modules

The final modules produced from each single algorithm and the consensus were evaluated using Pascal[18] (Pathway scoring algorithm). Pascal implements a fast and rigorous gene scoring and pathway enrichment pipeline that can be run on a local machine. The SNP values are converted to gene scores by computing pairwise SNP-by-SNP correlations and obtaining Z-scores from their distribution. These obtained gene scores are fused with the pathway enrichment analysis to recompute a chi-square P-value for the given set of module genes. Thus, the obtained chi-square P-value serves as the significance of the module in its enrichment of the disease-associated pathway gene loci. A combined P-value was computed for each of the methods using Fisher’s method[19], diseases, and datasets for ranking the performance of the modules in each criterion.

### Integration of MS single-omic modules

Clique SuM was ranked as the best performing method on average for both transcriptomic and methylomic data, according to the MS GWAS enrichment of the modules calculated by Pascal. Therefore, significant Clique SuM modules (P < 0.05) were selected for further analysis (nine transcriptomic and four methylomic modules). Consensus modules were generated across each omic by applying a module count-based method, where the criteria for gene inclusion in the consensus is its presence in a certain number of single-method modules. To balance the weight of each omic in the multi-omics integration, the top four significant modules per omic were used to create each consensus (Fig. 4a, b). Single-omic Clique SuM consensus were ranked again by GWAS enrichment, and the best performing consensus per omic was selected for integration into the multi-omics module.

### Enrichment analyses of the MS multi-omics module

Disease enrichment analysis of the multi-omics module was performed by Fisher’s exact test, with a significance threshold of P < 0.05. MS-associated genes were obtained from the gene-disease association summary provided by DisGeNET database 6.0[20]⍰. All genes with a known association to the disease “multiple sclerosis” (Unified Medical Language System unique identifier C0026769) were considered MS-associated genes (n = 1,105). Pathway enrichment analysis was carried out using the function *enrichKEGG* from the clusterProfiler R package[21]⍰, version 3.14.3. P-values were adjusted for multiple testing using Benjamini-Hochberg FDR correction, with a significance threshold of adj. P < 0.05. Enrichment of the multi-omics module in MS risk-factor-associated genes was performed by Fisher’s exact test, with a significance threshold of P < 0.05. To provide a uniform comparison of MS risk factor-associated genes across datasets, the module was tested for enrichment in the top 1,000 DMGs (with at least P < 0.05) obtained from the differential methylation analysis with ChAMP for each risk factor dataset.

### Representation of the MS multi-omics module

Experimentally validated interactions for the multi-omics module genes were obtained from STRING database version 11 (experimental score > 700) and imported into Cytoscape[22] version 3.7.2. To determine representative functional clusters of module genes, overrepresented Gene Ontology (GO) Biological Process (BP) terms in the module were found using BiNGO[23] version 3.0.4, with Benjamini-Hochberg FDR for multiple testing correction, and a significance threshold of adj. P < 0.05. Then, enriched GO terms with adj. P < 1×10^−10^ were summarized using REVIGO[24] server tool (medium allowed similarity = 0.7) and categories of interest were selected by uniqueness (>= 80 %), dispensability (>= 50 %), and frequency (<= 10 %) criteria. Further manual assessment was performed to group similar terms with an adequate number of genes in the network.

## RESULTS

### A benchmark comparing 337 transcriptionally derived disease modules from 19 different diseases

We compiled a benchmark source of disease modules and summary statistics of GWAS datasets from 19 well-powered case-control studies (Supplementary Table 1), some of which were previously used in the DREAM topological disease module challenge[12]. For these datasets we assessed modules using the same metric as in the recent DREAM study[12], based on the pathway scoring algorithm (Pascal)[18]. For each disease we compiled one to five publicly available transcriptomic datasets considering both easily assessable tissues (e.g. blood) and target tissues, thereby covering 47 transcriptomic datasets in total (Fig. 1a). Modules were created using eight different methods from MODifieR[13]. In addition, we also tested if genes detected by several methods, hereafter called consensus module genes, had higher enrichment scores than single-method module genes. Enrichment scores for the non-empty modules (n = 337) from this analysis were summarized for each method and dataset (Fig. 2a). In total, we found significantly GWAS-enriched modules in 17.8% (60/337) of the single-method modules and 25.5% (12/47) of the non-empty consensus modules that combined at least three methods as a criterion. These numbers seemed higher than expected, which might have been a consequence of the same GWAS being used to evaluate multiple transcriptomic datasets of the same disease. Hence, we aggregated scores of the same disease and method as meta P-values (see Methods). Out of the 152 possible disease-method combinations, 18% of the pairs showed a significant GWAS Pascal enrichment, which is more than expected by chance (n = 27, P = 1.0 × 10^−8^). The most enriched method was Clique SuM, which showed significant enrichment in seven out of 19 diseases (binomial test P = 2.3 × 10^−5^). Many methods exhibited strong enrichments in coronary artery disease (CAD), type 2 diabetes, multiple sclerosis (MS), rheumatoid arthritis (RA), and the inflammatory bowel diseases(IBD), ulcerative colitis (UC) and Crohn’s disease (CD), while no significant enrichments were found for asthma, hepatitis C, type 1 diabetes, narcolepsy, Parkinson’s disease, or for any psychiatric and social diseases. If we instead ranked methods based on their respective module GWAS enrichment, Clique SuM showed significant association in 34% (16/47) of the modules corresponding to seven different diseases followed by consensus modules identified by two out of three methods. Lastly, DIAMOnD and co-expression-based methods all achieved significant results, although worse than Clique SuM.

**Figure 1.**
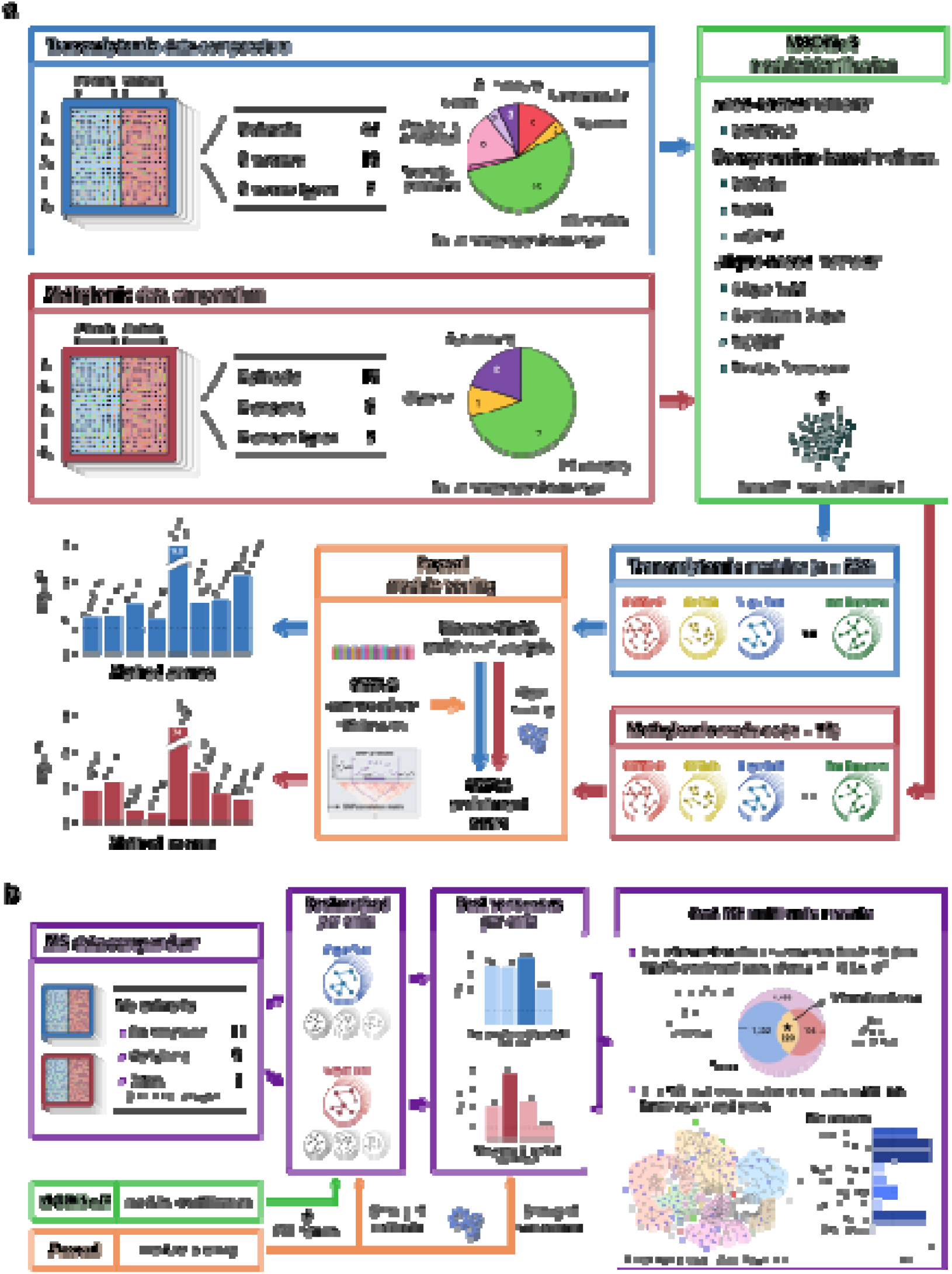
Overview of the benchmark assessment of disease modules and the integration workflow for MS. (**a**) Transcriptomic and methylomic datasets from 19 different diseases were used as inputs for eight MODifieR module identification methods. The resulting single-omic disease modules (n = 456) were independently assessed by GWAS enrichment analysis of the same disease using Pascal module scoring. MODifieR methods were evaluated by the combined enrichment score of their respective disease modules. (**b**) Multi-omic integrative workflow for multiple sclerosis (MS)-associated modules. Data from 20 case-control comparisons were used as input for module detection with MODifieR methods. Clique SuM modules presented the highest GWAS enrichment score and were therefore used to generate single-omic consensus modules. The intersection of the best transcriptomic and methylomic consensus modules resulted in an MS multi-omic module (n = 220 genes) with the highest GWAS enrichment, which was independently found to be enriched for genes associated with five known lifestyle MS risk factors using public omics data from healthy individuals.

**Figure 2.**
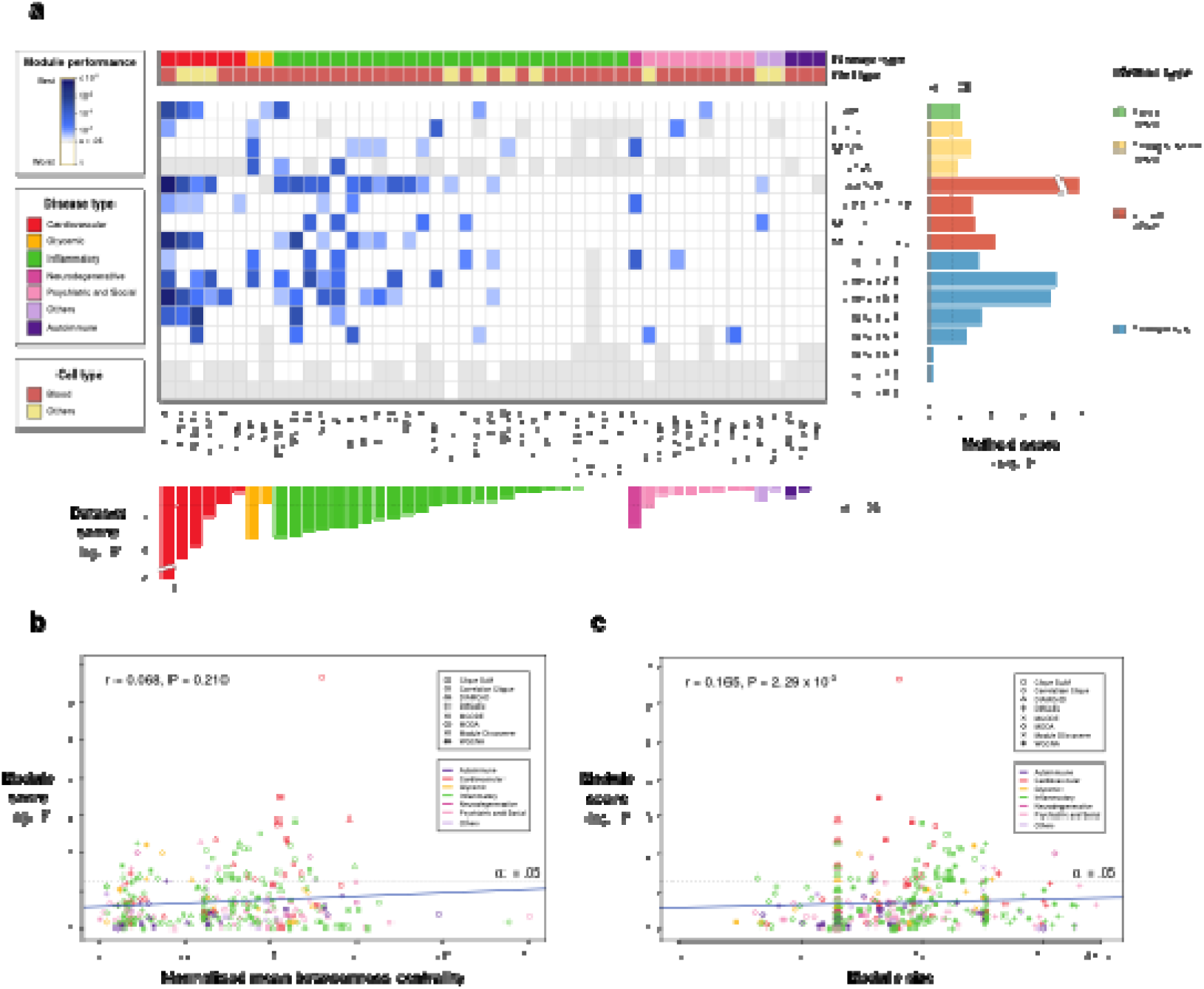
Genomic concordance of MODifieR modules on transcriptomic datasets. (**a**) Heatmap of PASCAL p-values for eight single-method and eight consensus MODifieR modules, identified for 47 publicly available transcriptomic datasets. Module performance P-values are shown in a white to blue scale, where any shade of blue represents a significant module (< 0.05; the darker, the more significant), white represents a non-significant module, and grey represents a module of size zero. Datasets are classified into six disease types: cardiovascular (red), glycemic (golden), inflammatory (green), neurodegenerative (fuchsia), psychiatric and social (pink), autoimmune (dark purple), and others (light purple); and two cell types: blood (maroon), and others (light yellow). Datasets are ranked by meta P-values using Fisher’s method of the single-method module P-values across and within their disease types (dataset score, bottom boxplot). MODifieR methods are organized by algorithm type: seed-based (green), co-expression-based (yellow), and clique-based (red), plus the consensus modules (blue). Single-methods and consensus were scored by meta P-values across datasets (method score, right boxplot). Consensus x/8 indicates that the module genes are found in at least x methods out of eight. (**b**) Scatter plot showing Spearman correlation between module score and betweenness centrality. Modules are represented with a different shape depending on their method and colored based on the disease type. (**c**) Scatter plot showing Spearman correlation between module score and module size. Modules are represented with a different shape depending on their method and colored based on the disease type.

Next, we tested the impact of network centrality and module size as potential confounding factors of the applied performance metric. We found a significant but very modest correlation for module size (Fig. 2c, Spearman rho = 0.165, P = 2.3 × 10^−3^), and a non-significant correlation for interactome centrality (Fig. 2b, rho = 0.068, P = 0.21). Thus, it is meaningful to compare results with differences in those module properties. In summary, we found that the Clique SuM method resulted in the highest disease enrichment for most diseases, while not producing significant modules for others, such as type 2 diabetes, where co-expression-based methods and DIAMOnD scored best. In general, we observed stronger enrichments for inflammatory diseases and weaker results for psychiatric and social diseases. Considering that the transcriptomic modules showed that Clique SuM was the best performing method and that the cardiovascular and inflammatory diseases were the most enriched within the Clique SuM modules, we wanted to test whether this was true for methylomic data as well.

### A benchmark comparing 72 methylation-based disease modules from six different diseases using GWAS

Following the same logic of the transcriptomic benchmark, we performed a similar benchmark study for methylation modules. We collected ten datasets from three different disease categories, including six complex diseases, and ran the eight MODifieR methods on them (Fig. 1a). In addition, we constructed consensus modules for each of the datasets. Modules were then tested for GWAS enrichment using Pascal. Inspecting the overall performance, we found nine single-method modules with a significant GWAS enrichment (9/72, 11.8%). Though this might be due to disease and cell type heterogeneity, the enrichment is more than expected by chance (P=9.6x 10^−3^). Interestingly, the inflammatory diseases such as MS and UC showed a more significant GWAS enrichment Considering that the evaluation of module performance by GWAS enrichment may be biased due to differences in module sizes and interactome centrality, we again assessed the correlation between these values. We found a significant correlation between GWAS enrichment and module size (Fig. 3c, rho = 0.235, P = 0.046) and a non-significant correlation between GWAS enrichment and interactome centrality (Fig. 3b, rho = 0.190, P = 0.109). We found that 12.5% of the disease-method combinations yielded significant GWAS enrichment, which is more than expected from an independent random selection of modules (Fisher’s exact test P = 0.031, n = 6). The highly enriched disease modules belong to MS, UC and CD. Two out of the six diseases showed significant GWAS enrichment by using the Clique SuM modules (P = 0.032). In summary, Clique SuM method resulted in a more significant GWAS enrichment for most diseases also for the methylomic benchmark.

**Figure 3.**
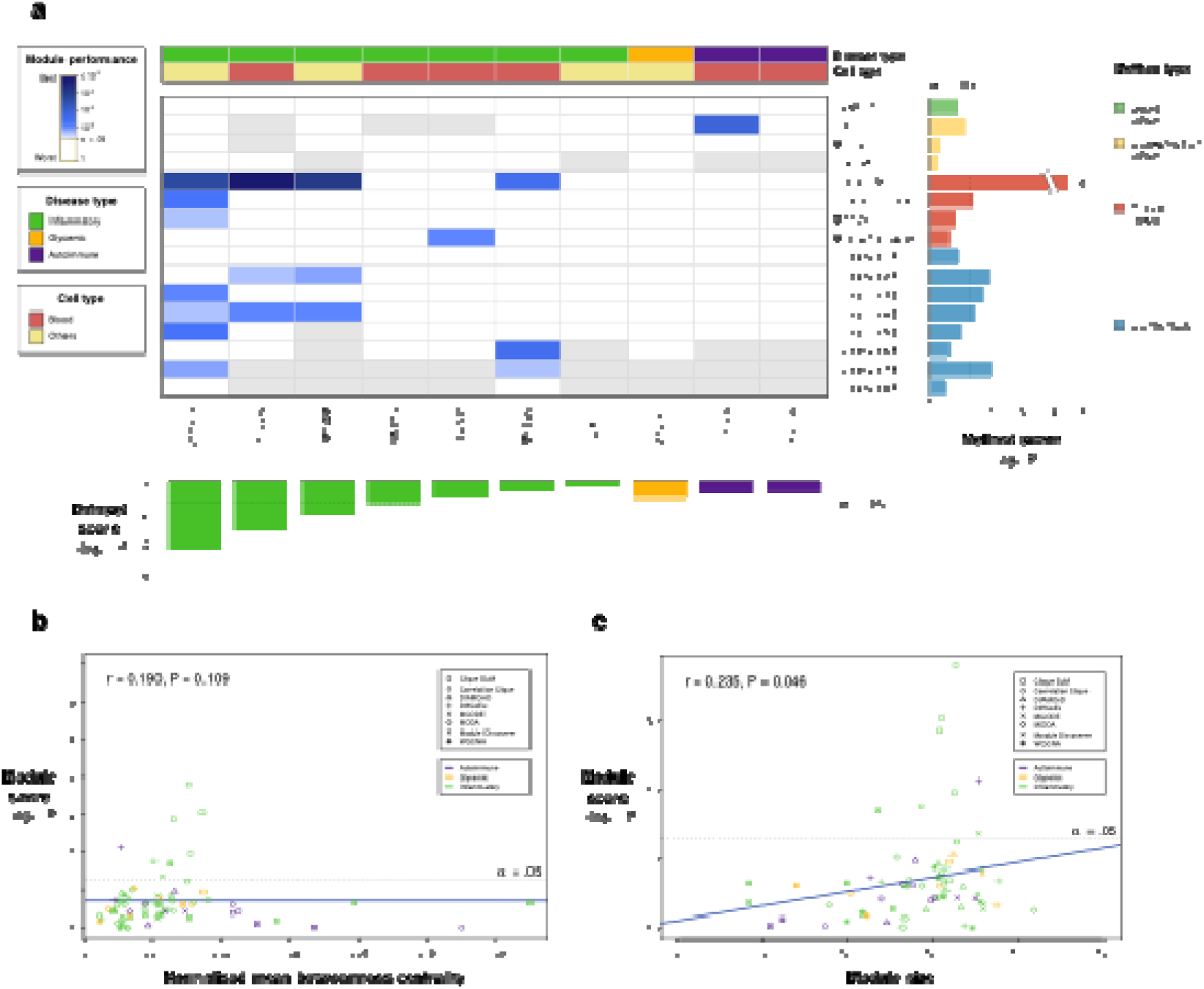
Genomic concordance of MODifieR modules on methylomic datasets. (**a**) Heatmap of Pascal p-values for eight single-method and eight consensus MODifieR modules, identified for ten publicly available methylomic datasets. Module performance P-values are shown in a white to blue scale, where any shade of blue represents a significant module (P < 0.05; the darker, the more significant), white represents a non-significant module, and grey represents a module of size zero. Datasets are classified into two disease types: glycemic (golden), and inflammatory (green); and two cell types: blood (maroon), and others (light yellow). Datasets are ranked by Fisher’s combined P of the single-method module P-values across and within their disease types (dataset score, bottom boxplot). MODifieR methods are organized by algorithm type: seed-based (green), co-expression-based (yellow), and clique-based (red), plus the consensus modules (blue). Single-methods and consensus are scored by meta P-values across datasets (method score, right boxplot). Consensus x/8 indicates that the module genes are found in at least x methods out of eight. (**b**) Scatter plot showing Spearman correlation between module score and betweenness centrality. Modules are represented with a different shape depending on their method and colored based on the disease type. (c) Scatter plot showing Spearman correlation between module score and module size. Modules are represented with a different shape depending on their method and colored based on the disease type.

**Figure 4.**
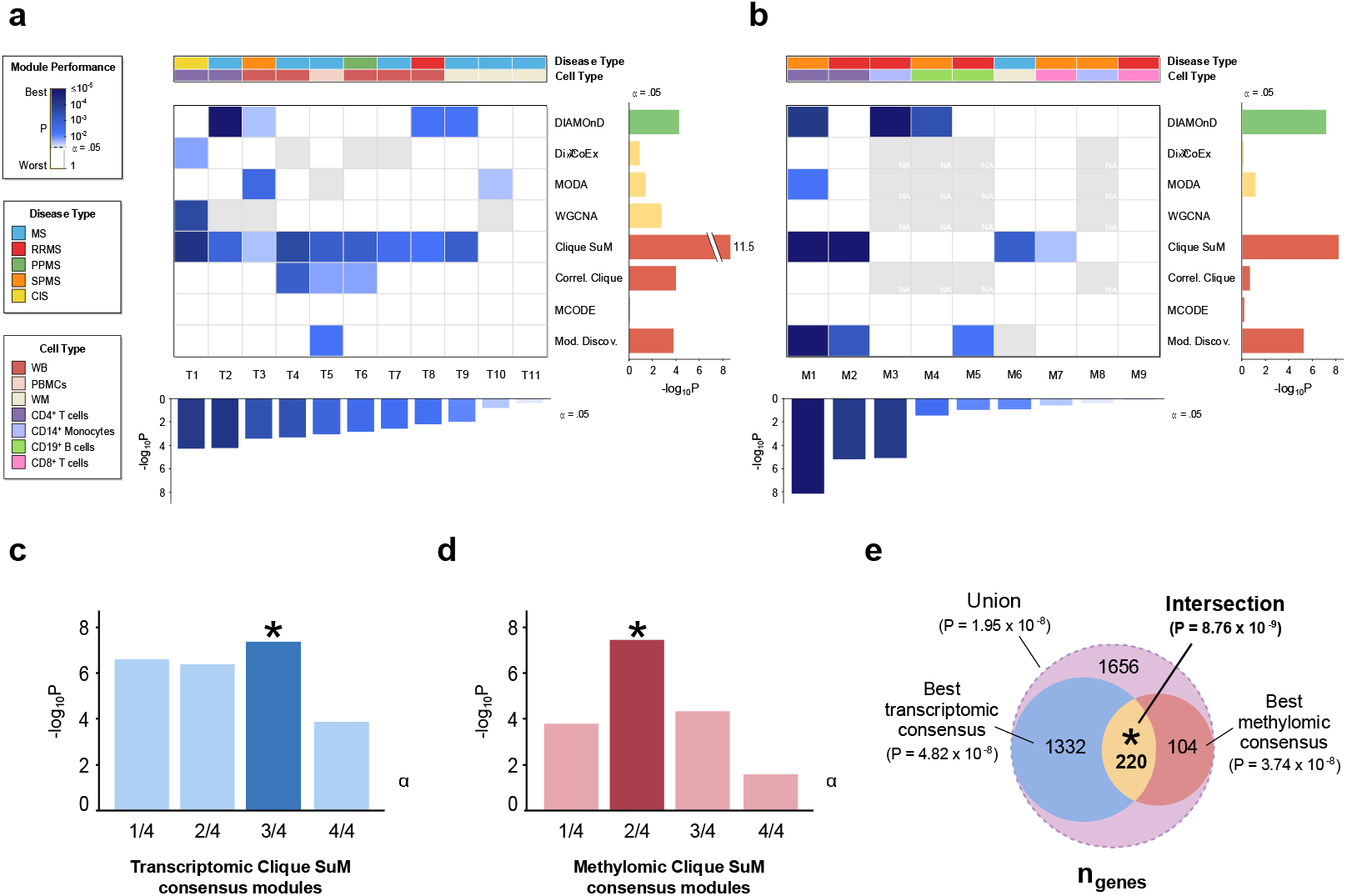
Genomic concordance of MODifieR modules on MS use case data. (**a**) Heatmap of PASCAL p-values for eight single-method MODifieR modules, identified for ten MS-related transcriptomic datasets. Module performance P-values are shown in a white to blue scale, where any shade of blue represents a significant module (P < 0.05), white represents a non-significant module, and grey represents a module of size zero. Datasets are classified into the reported MS type: MS (blue), RRMS (red), PPMS (green), SPMS (orange), and CIS (yellow); and four cell types: whole blood (maroon), PBMCs (light brown), white matter (light yellow), and CD4+ T cells (purple). Datasets are meta P-values of the single-method enrichments (dataset score, bottom boxplot). MODifieR methods are organized by algorithm type: seed-based (green), co-expression-based (yellow), and clique-based (red). Single methods are scored by P of the significant modules across datasets (method score, right boxplot). (**b**) Heatmap of PASCAL p-values for four single-method MODifieR modules, identified for nine MS-related transcriptomic datasets. (**c-d**) Bar plots of Pascal p-values for the MS consensus modules generated with Clique SuM from transcriptomic (a) and methylomic (b) datasets. (**e**) Union and intersection of the top performing modules, shown as a Venn diagram.

### Multi-omics approach revealed a module enriched for MS-associated genes

Considering genomic concordance as the guidance principle for the modules that show enrichment for GWAS SNPs, differentially methylated genes and differentially expressed genes, we further wanted to evaluate multiple datasets of one specific disease, i.e., MS. We compiled 11 MS transcriptomic datasets and nine methylation (Supplementary Table 2) comparisons from GEO which satisfy the pre-defined dataset criteria (see Methods). For each dataset we implemented the pipeline for module identification and scoring shown in Fig. 1b. We evaluated each module using MS SNP enrichment analysis and selected the most enriched modules per omic from this metric. This analysis again showed that Clique SuM yielded the far highest average enrichment score (meta P = 3.2 × 10^−12^) and was significantly enriched (P < 0.05) in 9/11 transcriptomic datasets (Fig. 4a) and 4/9 of the methylation datasets (Fig. 4b). From the significant modules generated by Clique SuM, we choose the top four modules from each of the gene transcription and methylation sets, and prioritized genes detected in modules from multiple datasets in each omic. This analysis showed that the strongest MS SNP enrichment was found for genes in at least three out of four transcriptomic modules (n=1,552; P= 6.0 × 10^−7^) and two out of four methylomic modules (n=324, P= 1.5×10^−6^). Next, we used the same principle to combine these two and found that the intersection between the gene transcription and methylation consensus resulted in a module (n = 220 genes, Fig. 4) enriched for MS-associated genes (75/220, P < 2.2 × 10^−16^, OR = 7.8) and with the highest GWAS enrichment (P = 8.8 × 10^−9^) which we hereafter referred to as the multi-omics MS module.

### The multi-omics MS module was enriched in genes associated with major MS pathways

As we used GWAS enrichment as a selection criterion, the high GWAS enrichment of the final module was partly expected, which led us to analyze its biological functions and their potential epigenetic associations to MS. First, pathway enrichment analysis showed that the multi-omics module genes are significantly involved in several inter-linked immune-related pathways, most of which have been previously associated to MS, including the T cell receptor[25] (adjusted P = 3.6 × 10^−47^), PI3K/Akt[26] (P = 4.6 × 10^−35^), ErbB[27] (P = 7.7 × 10^−32^), Fc epsilon RI[28] (P = 8.3 × 10^−30^), chemokine[29,30] (P = 2.6 × 10^−28^), MAPK[31,32] (P = 2.0 × 10^−25^), and B cell receptor[32] (P = 3.9 × 10^−19^) signaling pathways; Th17 (P = 9.6 × 10^−29^), and Th1 and Th2 (P = 6.9 × 10^−19^) cell differentiation[33]; natural killer cell mediated cytotoxicity (P = 1.6 × 10^−27^); and leukocyte transendothelial migration (P = 3.9 × 10^−20^), which indeed supports their relevance in MS. Interestingly, the module was also highly enriched in morphogenetic and neurogenetic signaling pathways, such as the neurotrophin (adjusted P = 1.3 × 10^−36^), Ras (P = 1.4 × 10^−36^), Rap1 (P = 2.2 × 10^−35^), vascular endothelial growth factor (VEGF, P = 1.7 × 10^−27^), FoxO (P = 3.6 × 10^−27^), and mTOR (P = 4.1 × 10^−14^) signaling pathways; and in growth hormone synthesis, secretion and action (P = 6.6 × 10^−31^).

### The multi-omics MS module was enriched in genes associated with five known environmental MS risk factors validated in an independent cohort

Second, from a literature study[34,35] we found nine environmental MS risk factors of varying evidence for which we could identify methylation studies in healthy controls. For each of these risk factors we derived the top 1000 differentially methylated genes (DMGs) and tested their enrichment with the module. Intriguingly, the module was significantly enriched for genes associated with five risk factors (Fig. 5b), which included the top associated risk factors, i.e., Epstein-Barr virus (EBV) infection (Fisher exact test P = 1.5 × 10^−3^, OR = 2.1) and smoking (P = 1.2 × 10^−4^, OR = 2.3), as well as low sun exposure (P = 1.2 × 10^−4^, OR = 2.3), high BMI (P = 0.023, OR = 1.7) and alcohol consumption (P = 2.9 × 10^−4^, OR = 2.2). Then, we asked whether these putative gene-risk factor associations could be validated using an independent omics dataset with paired risk factor associations. For this purpose, we utilized methylation arrays of peripheral blood from 139 MS patients and 140 controls, which have been described previously[36]. In this analysis we also considered risk factor associations for each individual including age, sex, BMI at age of 20, smoking, alcohol consumption, sun exposure, night shift work, contact with organic solvents. This enabled analysis of DMGs for the MS and risk factor status as covariates in linear mixed effect analysis. Indeed, the module genes were highly significantly enriched for MS (n = 217; permutation test P = 1.2 × 10^−47^), but also for all the tested risk factors (EBV was not included, Methods) and non-significantly associated to age and sex having 104-135 of the genes in each factor (3.9×10^−8^ < P < 0.013; Fig 5b). Combining all these results we found 90 of the 220 module genes to be associated with a risk factors from both the risk factor studies, 25 genes were associated with two risk factors, and seven genes were associated with three risk factors (CSK, PRKCA, PRKCZ, RUNX1, RUNX3, STAT5A, and SYNJ2) (Fig. 5c). These associations suggest that the multi-omics module is capturing a key disease network with both genetically and epigenetically driven alterations, thereby providing the possibility to use it to identify potential novel biomarkers or therapeutic targets for MS.⍰

**Figure 5.**
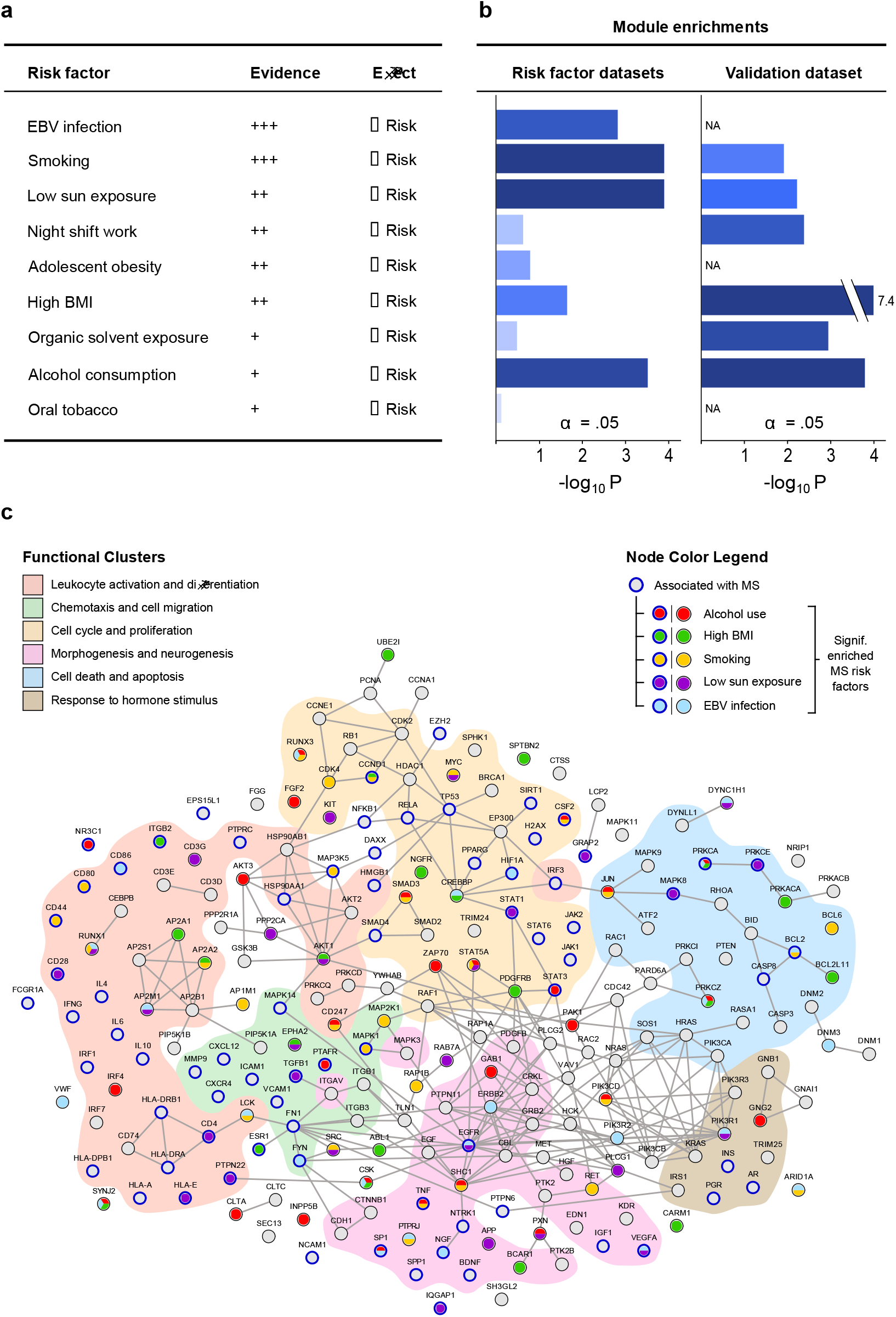
Risk factor enrichment and network visualization of the MS multi-omic module. (**a**) Evidence levels and effect on MS of the risk factor. ⍰ (**b**) Enrichment overlap of multi-omic MS module genes in the top 1,000 DMGs in risk factor datasets and independent risk factor methylation dataset (see Methods) shown as Fisher exact test P-values (threshold α=0.05). (**c**) Visualization of the module. Nodes (module genes) are arranged in functional clusters according to their overrepresented GO terms. Genes with a known association to MS are marked with a blue circle. Node colors display the associations to an MS risk factor for which the module is significantly enriched (red, alcohol use; green, high BMI; yellow, smoking; purple, low sun exposure; light blue, EBV infection; grey, no association). Edges were extracted from the STRINGdb v11 human PPI network of experimentally validated interactions (confidence score > 700).

## DISCUSSION

The analysis of case control data in the context of networks has gained increased interest to detect consistent robust gene signatures of individual diseases. The application of disease modules might vary for different researchers, but here we systematically aimed at the detection of disease genes supported by genetic association. For this purpose, our study of the transcriptome and methylome profiles of 19 diseases showed significant GWAS enrichments for several inflammatory and heart diseases, while psychiatric disorders showed no enrichments and might not be suitable for GWAS validation of modules, potentially due to differences in affected tissue types and sampling points. However, analysis of the significant results showed that methods based of differentially expressed cliques in the protein-protein interaction network demonstrated the strongest enrichments (highest scoring for Clique SuM), while those based primarily on correlations, like WGCNA, showed weak enrichments. A potential reason for this could be that GWAS has shown to be mostly associated to the central genes of the protein-protein interaction (PPI) network, but our analysis demonstrated that the correlation between GWAS enrichment and centrality was non-significant. We also tested whether there was an improvement using consensus approaches that counted the frequency of the result of multiple methods but found this not to increase performance. Moreover, we tested the same strategy on a set of inflammatory, glycemic, and autoimmune methylation datasets and found similar results. We would like to emphasize that, rather than scoring a single best working method, our result is a pipeline for evaluating modules using independent high-throughput enrichments.

The work on transcription and methylation datasets suggested that MS is a disease highly enriched for GWAS, and we therefore tested if increased enrichments could be derived by their integration. We found 20 publicly available datasets and run assessment for both omics independently, which again showed Clique SuM to score highest. We then tested if improved results could be obtained using modules from multiple datasets of these two omics using consensus modules from Clique SuM. This resulted in a module of 220 genes highly enriched for GWAS (P = 8.8 × 10^−9^). The multi-omic module was highly enriched in immune-associated pathways, such as T cell and B cell receptor signaling, Th1/Th2 differentiation, or leukocyte transendothelial migration. These results conform with the current hypothesis that MS is mediated by an autoreactive response of CD4+ T cells against myelin surrounding neuronal axons, preceded by their migration across the blood-brain barrier (BBB)[37]. This autoproliferation of brain-targeting Th1 cells has been shown to be driven by memory B cells, in a process mediated by HLA-DR15[38]. In addition, another enriched pathway was VEGF signaling. MS patients present high serum VEGF levels, which is related to pro-inflammatory functions and can alter the permeability of the BBB[39]. As GWAS was used for method prioritization we asked if modules instead could be validated using epigenetics and lifestyle risk factor genes that we identified to associate with MS. With this aim, we compiled a set of publicly available data from omics studies of these risk factors in healthy individuals. This analysis demonstrated that five out of eight risk factors were enriched in our module. In order to validate the use of an environmental assessment using public domain risk factor association we found an independent methylome study of MS comprising environmental data for each MS and healthy individual. This analysis showed a remarkable enrichment of the 220 module genes by 217 to differentially methylated genes for MS (P = 1.2 × 10^−47^), and a majority to be associated with the tested risk factors.

In contrast to previously known community challenges, in our study we not only used the topological property of the network, but we also combined the methods to use an omics-based input to uncover the disease modules that might be dysregulated at each omics level, contributing to the diverse causative mechanisms behind complex diseases. Although using the PPI network as background may lead to certain knowledge bias, this kind of benchmark allowed us to look at the relevant risk factors. In our assessment of the disease modules, methods such as Clique SuM and DIAMOnD did perform better than the community-based consensus predictions.

In summary, our study provides a practical integrative workflow that enables system-level analysis of heterogeneous diseases, in terms of multi-omics disease modules, as well as the validation of these by using both disease-specific GWAS and risk factors enrichment. We believe that this analysis validates our integrated use datasets and suggest a pipeline that readily could be tested in at least in other autoimmune and cardiovascular diseases. Lastly, our study did not aim to optimize hyper-parameters for individual disease modules, and instead used default values when possible, and to the methods from the MODifieR R package implementation of the methods[13]. However, this might be an important task for specific disease and our code and processed datasets are available at GitLab (https://gitlab.com/Gustafsson-lab/modifier-benchmark). In future work, this approach can be expanded to include diverse and context-specific networks to determine whether our multi-omics modules are able to capture various other levels of granularity.

## Supporting information

All case-control comparisons used in the Transcriptomic and Methylomic benchmarks.

All case-control comparisons used in the MS use case benchmark.

All Methods implemented in the benchmark.

## DECLARATIONS

## ETHICS APPROVAL AND CONSENT TO PARTICIPATE

Not applicable

## AVAILABILITY OF DATA AND MATERIALS

The data used for transcriptomic benchmark and methylation benchmark are downloaded from GEO. The disease specific GWAS files are downloaded from the latest Pascal version. The processed Data for analysis is available at https://gitlab.com/Gustafsson-lab/modifier-benchmark.The risk factor (EIMS) data will be made available on request. The R-package MODifieR is available on the GitLab: https://gitlab.com/Gustafsson-lab/MODifieR; the code used for benchmark analysis and risk factor analysis is available on GitLab: https://gitlab.com/Gustafsson-lab/modifier-benchmark; the latest Pascal version: https://www2.unil.ch/cbg/index.php?title=Pascal.

## COMPETING INTERESTS

The authors declare no competing interests.

## FUNDING

This work was supported by the Swedish Research Council (grant 2015-03807(M.G.), grant 2018-02638(M.J.)), the Swedish foundation for strategic research (grant SB16-0095(M.G.)), the Center for Industrial IT (CENIIT)(M.G.), European Union Horizon 2020/European Research Council Consolidator grant (Epi4MS, grant 818170(M.J.)), Knut and Alice Wallenberg Foundation (grant 2019.0089(M.J.)) and the Knowledge Foundation (grant 20170298 (Z.L.)). Computational resources were granted by Swedish National Infrastructure for Computing (SNIC; SNIC 2020/5-177, LiU-2018-12 and LiU-2019-25).

## AUTHOR CONTRIBUTIONS

T.V.S.B. compiled the necessary data for the benchmark analysis. H.A.W. performed the transcriptomic benchmark analysis. T.V.S.B. performed the methylation benchmark analysis. D.M.E. and H.A.W. performed the MS use case analysis. D.M.E performed the risk factor analysis. M.J.,I.K.,T.O., and L.A., provided the raw data and collected the associated risk factor data for the independent methylation dataset. T.V.S.B performed the independent validation dataset analysis. T.V.S.B. and D.M.E. collectively made the plots and figures for the manuscript. M.G. and Z.L. designed the study. T.V.S.B. and D.M.E. prepared the manuscript. All authors discussed the results and commented on the manuscript at all stages.

## SUPPLEMENTARY MATERIALS

Supplementary Table 1: All case-control comparisons used in the Transcriptomic and Methylomic benchmarks.

Supplementary Table 2: All case-control comparisons used in the MS use case benchmark.

Supplementary Table 3: All Methods implemented in the benchmark.

## Notes

https://gitlab.com/Gustafsson-lab/modifier-benchmark.

### Competing Interest Statement

The authors have declared no competing interest.

### Summary of Updates

The title, abstract were revised

