## Supplementary material for "A validated generally applicable approach using the systematic assessment of disease modules by GWAS reveals a multi-omic module strongly associated with risk factors in multiple sclerosis": All case-control comparisons used in the Transcriptomic and Methylomic benchmarks.

**Supplementary Table 1 |** All case-control comparisons used in the Transcriptomic and Methylomic benchmarks.

**Transcriptomic Benchmark case-control comparisons**

| Disease Type | Disease | celltype | Dataset | PMID | Platform | controls | patients | gwas | gwas_reference | download | Label (Fig. 2a) |
|---|---|---|---|---|---|---|---|---|---|---|---|
| Psychiatric and Social | Anorexia | DLPFC | GSE60190 | 25180571 | GPL6947 | 102 | 15 | EUR.GCAN.Anorexia.gcan_meta.out.txt | Boraska et al., Mol Psychiatry (2014) | http://www.med.unc.edu/pgc/downloads | ANO DLPFC |
| Psychiatric and Social | Anxiety | wholeblood | GSE61672 | 25300922 | GPL10558 | 179 | 157 | anxiety.meta.full.cc.tbl.txt | Otowa et al., Mol Psychiatry (2016) | https://www.med.unc.edu/pgc/results-and-downloads | ANX WB |
| Inflammatory | Asthma | wholeblood | GSE69683 | 27925796 | GPL13158 | 87 | 334 | input_gabriel_pran.txt | Moffat et al., NEJM (2010) | https://www.cnrgh.fr/gabriel/results.html | AST WB |
| Inflammatory | Asthma | sputum | GSE76262 | 28179442 | GPL13158 | 21 | 93 | input_gabriel_pran.txt | Moffat et al., NEJM (2010) | https://www.cnrgh.fr/gabriel/results.html | AST Sputum |
| Inflammatory | Asthma | bronchialbiopsy | GSE76225 | NA | GPL13158 | 35 | 56 | input_gabriel_pran.txt | Moffat et al., NEJM (2010) | https://www.cnrgh.fr/gabriel/results.html | AST BroBio |
| Inflammatory | Asthma | epithelialbrusing | GSE76226 | NA | GPL13158 | 36 | 63 | input_gabriel_pran.txt | Moffat et al., NEJM (2010) | https://www.cnrgh.fr/gabriel/results.html | AST EpiBru |
| Psychiatric and Social | Autism | PBMC | GSE18123 | 23227143 | GPL570 | 33 | 31 | EUR.onqah.pgcasdeuro.txt.gz | PGC, Lancet (2013) | https://www.med.unc.edu/pgc/results-and-downloads | AUT 1 PBMC |
| Psychiatric and Social | Autism | PBMC | GSE18123 | 23227143 | GPL6244 | 82 | 41 | EUR.onqah.pgcasdeuro.txt.gz | PGC, Lancet (2013) | https://www.med.unc.edu/pgc/results-and-downloads | AUT 2 PBMC |
| Psychiatric and Social | Bipolar_Disorder | PBMC | GSE39653 | 23064081 | GPL10558 | 24 | 8 | BPSCZ.bp-only.results.txt | Ruderfer et al., Mol Psychiatry (2014) | https://www.med.unc.edu/pgc/results-and-downloads | BPD 1 PBMC |
| Psychiatric and Social | Bipolar_Disorder | PBMC | GSE18312 | 19582768 | GPL5175 | 8 | 9 | BPSCZ.bp-only.results.txt | Ruderfer et al., Mol Psychiatry (2014) | https://www.med.unc.edu/pgc/results-and-downloads | BPD 2 PBMC |
| Cardiovascular | CAD | CD4 | GSE9820 | 19059264 | GPL6255 | 14 | 20 | EUR.ASN.CAD.cad.add.160614.website.txt | Nikpay et al., Nat Genet (2015) | http://www.cardiogramplusc4d.org/downloads/ | CAD CD4 |
| Cardiovascular | CAD | CD14 | GSE9821 | 19059264 | GPL6255 | 13 | 18 | EUR.ASN.CAD.cad.add.160614.website.txt | Nikpay et al., Nat Genet (2015) | http://www.cardiogramplusc4d.org/downloads/ | CAD CD14 |
| Cardiovascular | CAD | Macrophages | GSE9821 | 19059264 | GPL6255 | 15 | 19 | EUR.ASN.CAD.cad.add.160614.website.txt | Nikpay et al., Nat Genet (2015) | http://www.cardiogramplusc4d.org/downloads/ | CAD Macro |
| Cardiovascular | CAD | Stemcells | GSE9821 | 19059264 | GPL6255 | 11 | 12 | EUR.ASN.CAD.cad.add.160614.website.txt | Nikpay et al., Nat Genet (2015) | http://www.cardiogramplusc4d.org/downloads/ | CAD Stem |
| Cardiovascular | CAD | subcutaneous_adipose | GSE64554 | 26645979 | GPL6947 | 10 | 13 | EUR.ASN.CAD.cad.add.160614.website.txt | Nikpay et al., Nat Genet (2015) | http://www.cardiogramplusc4d.org/downloads/ | CAD SubAdi |
| Cardiovascular | CAD | epicardial_adipose | GSE64554 | 26645979 | GPL6947 | 10 | 13 | EUR.ASN.CAD.cad.add.160614.website.txt | Nikpay et al., Nat Genet (2015) | http://www.cardiogramplusc4d.org/downloads/ | CAD EpiAdi |
| Inflammatory | Crohns_disease | CD4 | GSE87650 | 27886173 | GPL10558 | 17 | 21 | EUR.IBDGenetics.CD.txt | Franke et al., Nat Genet (2010) | http://www.ibdgenetics.org/projects.html | CD CD4 |
| Inflammatory | Crohns_disease | CD8 | GSE87650 | 27886173 | GPL10558 | 15 | 20 | EUR.IBDGenetics.CD.txt | Franke et al., Nat Genet (2010) | http://www.ibdgenetics.org/projects.html | CD CD8 |
| Inflammatory | Crohns_disease | CD14 | GSE87650 | 27886173 | GPL10558 | 18 | 25 | EUR.IBDGenetics.CD.txt | Franke et al., Nat Genet (2010) | http://www.ibdgenetics.org/projects.html | CD CD14 |
| Inflammatory | Crohns_disease | wholeblood | GSE87650 | 27886173 | GPL10558 | 24 | 23 | EUR.IBDGenetics.CD.txt | Franke et al., Nat Genet (2010) | http://www.ibdgenetics.org/projects.html | CD WB |
| Psychiatric and Social | Depression | PBMC | GSE39653 | 23064081 | GPL10558 | 24 | 21 | Depression.txt | NA | NA | DEP PBMC |
| Others | Macular_Degeneration | RPE | GSE510195 | 24265543 | GPL17629 | 7 | 9 | EUR.Fritsche_2015_AdvancedAMD.txt | Fritsche et al., Nat Genet (2015) | http://csg.sph.umich.edu/abecasis/public/amd2015/ | MD RPE |
| Inflammatory | MS | CD4 | GSE13732 | 18689680 | GPL570 | 30 | 39 | EUR.WTCCC2_MS_official.txt | IMSGC. Nature (2011) | NA | MS CD4 |
| Inflammatory | MS | PBMC | GSE41848 | 23748426 | GPL16209 | 38 | 54 | EUR.WTCCC2_MS_official.txt | IMSGC. Nature (2011) | NA | MS 1 PBMC |
| Inflammatory | MS | PBMC | GSE41849 | 23748426 | GPL16209 | 22 | 21 | EUR.WTCCC2_MS_official.txt | IMSGC. Nature (2011) | NA | MS 2 PBMC |
| Autoimmune | Narcolepsy | wholeblood | GSE21592 | NA | GPL571 | 10 | 10 | EUR.Hor2010.Narcolepsy.META_NARCO_HLAall_IMPUTED.txt | Hor et al., Nat Genet (2010) | | NARC WB |
| Others | Pancreatic_cancer | pancreas | GSE15471 | 19260470 | GPL570 | 39 | 39 | EUR.phs000206.pha002889.pancreatic_cancer.txt | Li et al., Carcinogenesis (2012) | http://www.ncbi.nlm.nih.gov/projects/gap/cgi-bin/analysis.cgi?study_id=phs000206.v4.p3&pha=2889 | PC Pancreas |
| Neurodegenerative | Parkinsons_disease | wholeblood | GSE6613 | 17215369 | GPL96 | 22 | 50 | EUR.phs000089_PD_phs000089.pha002868.txt.gz | NA | NA | PD WB |
| Inflammatory | RA | CD4 | GSE4588 | NA | GPL570 | 10 | 8 | EUR.Okada2014.RA_GWASmeta_European_v2.txt | Okada et al., Nature (2014) | http://plaza.umin.ac.jp/~yokada/datasource/software.htm | RA 1 CD4 |
| Inflammatory | RA | B | GSE4588 | NA | GPL570 | 9 | 7 | EUR.Okada2014.RA_GWASmeta_European_v2.txt | Okada et al., Nature (2014) | http://plaza.umin.ac.jp/~yokada/datasource/software.htm | RA B |
| Inflammatory | RA | CD4 | GSE56649 | 25880754 | GPL570 | 9 | 13 | EUR.Okada2014.RA_GWASmeta_European_v2.txt | Okada et al., Nature (2014) | http://plaza.umin.ac.jp/~yokada/datasource/software.htm | RA 2 CD4 |
| Inflammatory | RA | CD14 | GSE57386 | 25333715 | GPL13158 | 19 | 9 | EUR.Okada2014.RA_GWASmeta_European_v2.txt | Okada et al., Nature (2014) | http://plaza.umin.ac.jp/~yokada/datasource/software.htm | RA CD14 |
| Psychiatric and Social | Schizophrenia | PBMC | GSE18312 | 19582768 | GPL5175 | 8 | 13 | EUR.pgc.scz2.2014.txt | PGC, Nature (2014) | http://www.med.unc.edu/pgc/downloads | SCZ PBMC |
| Autoimmune | T1D | PBMC | GSE9006 | 17595242 | GPL96 | 24 | 43 | EUR.phs000180_T1D_phs000180.pha002862.txt | Barrett et al., Nat Genet (2009) | NA | T1D 1 PBMC |
| Autoimmune | T1D | PBMC | GSE9006 | 17595242 | GPL97 | 24 | 43 | EUR.phs000180_T1D_phs000180.pha002862.txt | Barrett et al., Nat Genet (2009) | NA | T1D 2 PBMC |
| Glycemic | T2D | PBMC | GSE9006 | 17595242 | GPL96 | 24 | 12 | EUR.T2D.Transethnic.txt | NA | http://diagram-consortium.org/downloads.html | T2D 1 PBMC |
| Glycemic | T2D | PBMC | GSE9006 | 17595242 | GPL97 | 24 | 12 | EUR.T2D.Transethnic.txt | NA | http://diagram-consortium.org/downloads.html | T2D 2 PBMC |
| Inflammatory | Ulcerative_colitis | CD4 | GSE87651 | 27886173 | GPL10558 | 17 | 20 | EUR.IBDGenetics.UC.txt.gz | Anderson et al., Nat Genet (2011) | http://www.ibdgenetics.org/projects.html | UC CD4 |
| Inflammatory | Ulcerative_colitis | CD8 | GSE87652 | 27886173 | GPL10558 | 15 | 20 | EUR.IBDGenetics.UC.txt.gz | Anderson et al., Nat Genet (2011) | http://www.ibdgenetics.org/projects.html | UC CD8 |
| Inflammatory | Ulcerative_colitis | CD14 | GSE87653 | 27886173 | GPL10558 | 18 | 20 | EUR.IBDGenetics.UC.txt.gz | Anderson et al., Nat Genet (2011) | http://www.ibdgenetics.org/projects.html | UC CD14 |
| Inflammatory | Ulcerative_colitis | wholeblood | GSE87655 | 27886173 | GPL10558 | 24 | 22 | EUR.IBDGenetics.UC.txt.gz | Anderson et al., Nat Genet (2011) | http://www.ibdgenetics.org/projects.html | UC WB |
| Inflammatory | Hepatitis_C | CD4 | GSE49954 | 24130824 | GPL570 | 5 | 5 | EUR.Rauch2010.HepC.META_RESOLVER_IMPUTED.txt.gz | Rauch et al., Gastroenterology (2010 | NA | HEPC H CD4 |
| Inflammatory | Hepatitis_C | CD4 | GSE49954 | 24130824 | GPL570 | 5 | 5 | EUR.Rauch2010.HepC.META_RESOLVER_IMPUTED.txt.gz | Rauch et al., Gastroenterology (2010 | NA | HEPC L CD4 |
| Inflammatory | Hepatitis_C | CD8 | GSE49954 | 24130824 | GPL570 | 5 | 5 | EUR.Rauch2010.HepC.META_RESOLVER_IMPUTED.txt.gz | Rauch et al., Gastroenterology (2010 | NA | HEPC H CD8 |
| Inflammatory | Hepatitis_C | CD8 | GSE49954 | 24130824 | GPL570 | 5 | 5 | EUR.Rauch2010.HepC.META_RESOLVER_IMPUTED.txt.gz | Rauch et al., Gastroenterology (2010 | NA | HEPC L CD8 |
| Inflammatory | MS | wholeblood | GSE17048 | 20190274 | GPL6947 | 45 | 99 | EUR.WTCCC2_MS_official.txt | IMSGC. Nature (2011) | NA | MS WB |
| Inflammatory | MS | white matter | GSE126802 | 31023360 | GPL13497 | 9 | 18 | EUR.WTCCC2_MS_official.txt | IMSGC. Nature (2011) | NA | MS WM |

**Methylomic Benchmark case-control comparisons**

| Disease Type | Disease | celltype | Dataset | PMID | Platform | controls | patients | gwas | gwas_reference | download | Label (Fig. 3a) |
|---|---|---|---|---|---|---|---|---|---|---|---|
| Inflammatory | Ulcerative_colitis | Colon biopsy | GSE27899 | 22826509 | GPL8490 | 10 | 10 | EUR.IBDGenetics.UC.txt.gz | Anderson et al., Nat Genet (2011) | http://www.ibdgenetics.org/projects.html | UC Colon |
| Inflammatory | MS | PBMC | GSE106648 | 29921915 | GPL13154 | 140 | 139 | EUR.WTCCC2_MS_official.txt | IMSGC. Nature (2011) | | MS PBMC |
| Inflammatory | MS | white matter | GSE40360 | 24270187 | GPL13154 | 19 | 28 | EUR.WTCCC2_MS_official.txt | IMSGC. Nature (2011) | | MS WM |
| Inflammatory | RA | CD4 | GSE71841 | NA | GPL13154 | 12 | 12 | EUR.Okada2014.RA_GWASmeta_European_v2.txt | Okada et al., Nature (2014) | http://plaza.umin.ac.jp/~yokada/datasource/software.htm | RA CD4 |
| Inflammatory | Crohns disease | wholeblood | GSE105798 | 32375395 | GPL13154 | 3 | 8 | EUR.IBDGenetics.CD.txt | Franke et al., Nat Genet (2010) | http://www.ibdgenetics.org/projects.html | CD WB |
| Inflammatory | MS | CD4 | GSE130029 | 31427643 | GPL13154 | 11 | 20 | EUR.WTCCC2_MS_official.txt | IMSGC. Nature (2011) | | MS CD4 |
| Inflammatory | Crohns disease | Adipocyte Stem cells | GSE138311 | 32252817 | GPL23976 | 6 | 7 | EUR.IBDGenetics.CD.txt | Franke et al., Nat Genet (2010) | http://www.ibdgenetics.org/projects.html | CD ASC |
| Glycemic | T2D | Liver biopsy | GSE65057 | 26977391 | GPL13154 | 14 | 10 | EUR.T2D.Transethnic.txt | NA | http://diagram-consortium.org/downloads.html | T2D liver |
| Autoimmune | T1D | CD14 | GSE56606 | 21980303 | GPL8490 | 33 | 15 | EUR.phs000180_T1D_phs000180.pha002862.txt | Barrett et al., Nat Genet (2009) | NA | T1D CD14 |
| Autoimmune | T1D | CD4 | GSE56606 | 21980303 | GPL8490 | 35 | 17 | EUR.phs000180_T1D_phs000180.pha002862.txt | Barrett et al., Nat Genet (2009) | NA | T1D CD4 |
