## Supplementary material for "A validated generally applicable approach using the systematic assessment of disease modules by GWAS reveals a multi-omic module strongly associated with risk factors in multiple sclerosis": All case-control comparisons used in the MS use case benchmark.

**Supplementary Table 2 |** All case-control comparisons used in the MS use case benchmark.

**MS Transcriptomic Benchmark case-control comparisons**

| Disease type | Disease | Cell type | Dataset | PMID | Platform | Controls | Patients | GWAS | GWAS reference | Label (Fig. 4a) | Pascal P (Clique SuM) | Selected for multi-omics module |
|---|---|---|---|---|---|---|---|---|---|---|---|---|
| Inflammatory | MS | Whole blood | GSE17048 | 20190274 | GPL6947 | 45 | 99 | EUR.WTCCC2_MS_official.txt | IMSGC. Nature (2011) | T7 | 1,22E-02 | FALSE |
| Inflammatory | RRMS | Whole blood | GSE17048 | 20190274 | GPL6947 | 45 | 36 | EUR.WTCCC2_MS_official.txt | IMSGC. Nature (2011) | T8 | 1,69E-02 | FALSE |
| Inflammatory | SPMS | Whole blood | GSE17048 | 20190274 | GPL6947 | 45 | 20 | EUR.WTCCC2_MS_official.txt | IMSGC. Nature (2011) | T3 | 3,77E-02 | FALSE |
| Inflammatory | PPMS | Whole blood | GSE17048 | 20190274 | GPL6947 | 45 | 43 | EUR.WTCCC2_MS_official.txt | IMSGC. Nature (2011) | T6 | 4,53E-03 | TRUE |
| Inflammatory | MS | White matter (RIM) | GSE108000 | 29312322 | GPL13497 | 10 | 7 | EUR.WTCCC2_MS_official.txt | IMSGC. Nature (2011) | T11 | 1,47E-01 | FALSE |
| Inflammatory | MS | White matter (PL) | GSE108000 | 29312322 | GPL13497 | 10 | 7 | EUR.WTCCC2_MS_official.txt | IMSGC. Nature (2011) | T9 | 4,77E-03 | TRUE |
| Inflammatory | MS | Whole blood | GSE41848 | 23748426 | GPL16209 | 79 | 133 | EUR.WTCCC2_MS_official.txt | IMSGC. Nature (2011) | T4 | 1,29E-03 | TRUE |
| Inflammatory | MS | PBMC | GSE21942 | 22021740 | GPL570 | 15 | 14 | EUR.WTCCC2_MS_official.txt | IMSGC. Nature (2011) | T5 | 5,30E-03 | FALSE |
| Inflammatory | MS | CD4+ T cells (unstim.) | GSE78244 | 27626663 | GPL17077 | 14 | 14 | EUR.WTCCC2_MS_official.txt | IMSGC. Nature (2011) | T2 | 6,37E-03 | FALSE |
| Inflammatory | MS | White matter | GSE126802 | 31023360 | GPL13497 | 9 | 18 | EUR.WTCCC2_MS_official.txt | IMSGC. Nature (2011) | T10 | 1,85E-01 | FALSE |
| Inflammatory | MS (CIS) | CD4+ T cells | GSE13732 | 18689680 | GPL570 | 30 | 39 | EUR.WTCCC2_MS_official.txt | IMSGC. Nature (2011) | T1 | 4,42E-04 | TRUE |

**MS Methylomic Benchmark case-control comparisons**

| Disease type | Disease | Cell type | Dataset | PMID | Platform | Controls | Patients | GWAS | GWAS reference | Label (Fig. 4b) | Pascal P (Clique SuM) | Selected for multi-omics module |
|---|---|---|---|---|---|---|---|---|---|---|---|---|
| Inflammatory | SPMS | CD4+ T cells | GSE130029 | 31053557 | GPL13534 | 11 | 8 | EUR.WTCCC2_MS_official.txt | IMSGC. Nature (2011) | M1 | 2,23E-05 | TRUE |
| Inflammatory | RRMS | CD4+ T cells | GSE130029 | 31053557 | GPL13534 | 11 | 12 | EUR.WTCCC2_MS_official.txt | IMSGC. Nature (2011) | M2 | 9,48E-07 | TRUE |
| Inflammatory | SPMS | CD8+ T cells | GSE130030 | 31053557 | GPL13534 | 14 | 4 | EUR.WTCCC2_MS_official.txt | IMSGC. Nature (2011) | M7 | 4,82E-02 | TRUE |
| Inflammatory | RRMS | CD8+ T cells | GSE130030 | 31053557 | GPL13534 | 14 | 10 | EUR.WTCCC2_MS_official.txt | IMSGC. Nature (2011) | M9 | 1,22E-01 | FALSE |
| Inflammatory | RRMS | CD14+ monocytes | NA | 31053557 | GPL13534 | 13 | 10 | EUR.WTCCC2_MS_official.txt | IMSGC. Nature (2011) | M3 | 1,46E-01 | FALSE |
| Inflammatory | SPMS | CD14+ monocytes | NA | 31053557 | GPL13534 | 13 | 13 | EUR.WTCCC2_MS_official.txt | IMSGC. Nature (2011) | M8 | 6,32E-01 | FALSE |
| Inflammatory | MS | White matter | GSE40360 | 24270187 | GPL13534 | 19 | 28 | EUR.WTCCC2_MS_official.txt | IMSGC. Nature (2011) | M6 | 5,93E-03 | TRUE |
| Inflammatory | RRMS | CD19+ B cells | NA | 31053557 | GPL13534 | 10 | 12 | EUR.WTCCC2_MS_official.txt | IMSGC. Nature (2011) | M5 | 8,67E-01 | FALSE |
| Inflammatory | SPMS | CD19+ B cells | NA | 31053557 | GPL13534 | 10 | 5 | EUR.WTCCC2_MS_official.txt | IMSGC. Nature (2011) | M4 | 5,02E-01 | FALSE |

**Legend**

| | |
|---|---|
| RRMS | Relapsing-remitting MS |
| SPMS | Secondary progressive MS |
| PPMS | Primary progressive MS |
| CIS | Clinically isolated syndrome |
| RIM | Rim of chronic lesion |
| PL | Perilesion |
