## Supplementary material for "A validated generally applicable approach using the systematic assessment of disease modules by GWAS reveals a multi-omic module strongly associated with risk factors in multiple sclerosis": All Methods implemented in the benchmark.

Supplementary Table 3 | All Methods implemented in the benchmark.

| **Method** | **type** | **Uses PPI/network constructed from data** | **Uses Pathway /genesets For preprocessing** | **Produces modules of significance** | **Can be run on local machine** |
|---|---|---|---|---|---|
| DIAMonD | Random Walker | X | - | X | X |
| MCODE | Clique | X | - | X | X |
| Clique Sum | Clique | X | - | X | X |
| Clique Correlation | Clique | X | - | X | X |
| Module Discoverer | Clique | X | - | X | X |
| WGCNA | Co-expression | X | - | X | X |
| DiffCoEx | Co-expression | X | - | X | X |
| MODA | Co-expression | X | - | X | X |
